# Regulation of pancreatic fibrosis by acinar cell-derived exosomal miR-130a-3p via targeting of stellate cell PPAR-γ

**DOI:** 10.1101/2020.12.01.407585

**Authors:** Qiang Wang, Hao Wang, Qingxu Jing, Yang Yang, Dongbo Xue, Chen jun Hao, Weihui Zhang

**Affiliations:** Department of General Surgery,Laboratory of Hepatosplenic Surgery, Ministry of Education,The First Affiliated Hospital of Harbin Medical University, Harbin, Heilongjiang Province,China

**Author notes:** These authors are regarded as co-first authors (Qiang Wang, Hao Wang). Correspondingauthor:Dongbo Xue,Department of General Surgery, The First A ffiliated Hospital of Harbin Medical University, Youzheng Street 23, Harbin, C hina. 150001,. Chenjun Hao,Department of General Surgery, The First Affiliated Hospital of Harbin Medical University, Youzheng Street 23, Harbin, China. 150001,haochenj.

**Keywords:** exosomes, miR-130a-3p, pancreatitis fibrosis, PPAR-γ

## Abstract

As endogenous miRNA carriers,exosomes play a role in the pathophysiological process of various diseases. However, their functions and regulation mechanisms in pancreatic fibrosis remain unclear. In this study, an RNA microarray was used to detect differentially expressed exosomal miR-130a-3p in AR42J cells after taurolithocholate (TLC) treatment. mRNA-seq was used to screen the differentially expressed PPAR-γ after pancreatic stellate cell (PSC) activation. Fluorescence labeling of exosomes and dynamic tracing showed that exosomes can fuse with the cell membrane of PSCs and transport miR-130a-3p into PSCs. A luciferase reporter gene assay was used to confirm that miR-130a can bind to PPAR-γ to inhibit PPAR-γ expression. In vitro and in vivo functional experiments were performed for gain-of-function studies and loss-of-function studies, respectively. These studies showed that acinar cell-derived exosomal miR-130a-3p promotes PSC activation and collagen formation through targeting of cellular PPAR-γ. Knockdown of miR-130a-3p significantly improved pancreatic fibrosis. Notably, miR-130a-3p knockdown reduced serum levels of hyaluronic acid (HA) and β-amylase and increased the C-peptide to protect endocrine and exocrine pancreatic functions and the function of endothelial cells. The exosomal miR-130a-3p/PPAR-γ axis participates in the activation of PSCs and the mechanism of CP fibrosis, thus providing a potential new target for the treatment of chronic pancreatitis fibrosis.

## INTRODUCTION

Chronic pancreatitis (CP) is a common digestive disease in the clinic. Typical CP pathological manifestations include pancreatic parenchymal fibrosis, acinar cell atrophy, pancreatic duct deformation, inflammatory cell infiltration and extracellular matrix (ECM) deposition. The essence of fibrosis is the increase in ECM synthesis and relative decrease in ECM degradation. The compromised dynamic balance between synthesis and degradation results in excessive ECM deposition. Activation of pancreatic stellate cells (PSCs) is the most important factor in the process of pancreatic fibrosis. PSC activation can produce type I and type III collagen, fibronectin and laminin^[1]^. In normal pancreatic tissues, PSCs remain in a quiescent state and do not promote pancreatic fibrosis. However, various stimulating factors in pancreatic injury can activate transformation of PSCs to myofibroblasts, which express α-SMA (α-smooth muscle actin) and secrete a large amount of ECM^[2]^. Many studies have shown an increased number of PSCs in the pancreatic fibrosis area in human chronic pancreatitis specimens and animal pancreatic fibrosis models and positive α-SMA staining (PSC activation state) and collagen staining in double-stained sections, indicating that activated PSCs in the pancreas are an important source of fibrotic collagen production and play an important role in the development of pancreatic fibrosis^[3]^.

Pancreatic fibrosis is an important factor in the occurrence of pancreatic cancer. In most cases, CP is associated with pancreatic fibrosis occurrence. Thus, understanding the mechanism of CP fibrosis is of great significance to pancreatic cancer prevention. Siech et al^[4].^ demonstrated that acinar cells can promote PSC activation in an acute pancreatitis (AP) model induced by alcohol and fat. Patel et al. ^[5]^found that damaged acinar cells in CP accelerated the process of pancreatic fibrosis by promoting PSC activation. In this regard, what is the mechanism by which pancreatic acinar cells activate PSCs? We investigated the mechanism by which pancreatic acinar cells activate PSCs. In recent years, studies have shown that communication between cells is dependent on extracellular vesicles. Therefore, we must pay attention to the effect of extracellular vesicles on CP fibrosis. Our studies indicate that during CP occurrence, exosomes secreted by acinar cells have a high concentration of miRNAs^[6,7].^ Among them, miR-130a-3p shows significant expression, and the uptake of exosomes carrying miR-130a-3p activates PSCs to release large amounts of collagen fibers, causing fibrosis of the pancreas.

## RESULTS

### Differentially expressed miRNAs in exosomes

The microarray analysis of miRNAs showed that the exosomes released by AR42J cells contained multiple miRNAs, consistent with findings in previous reports (Figure 1). Compared with the cells in the control group, activated AR42J cells secreted exosomes containing 53 underexpressed miRNAs and 62 overexpressed miRNAs. These differentially expressed miRNAs may be communication messengers between cells. In the current study, the expression of miR-130a-3p was significantly increased (Figure 1B). Literature review indicated that miR-130a-3p is involved in fibrosis regulation, but its role in CP fibrosis remains unclear. Therefore, we performed further research on miR-130a-3p.

**Figure 1.**
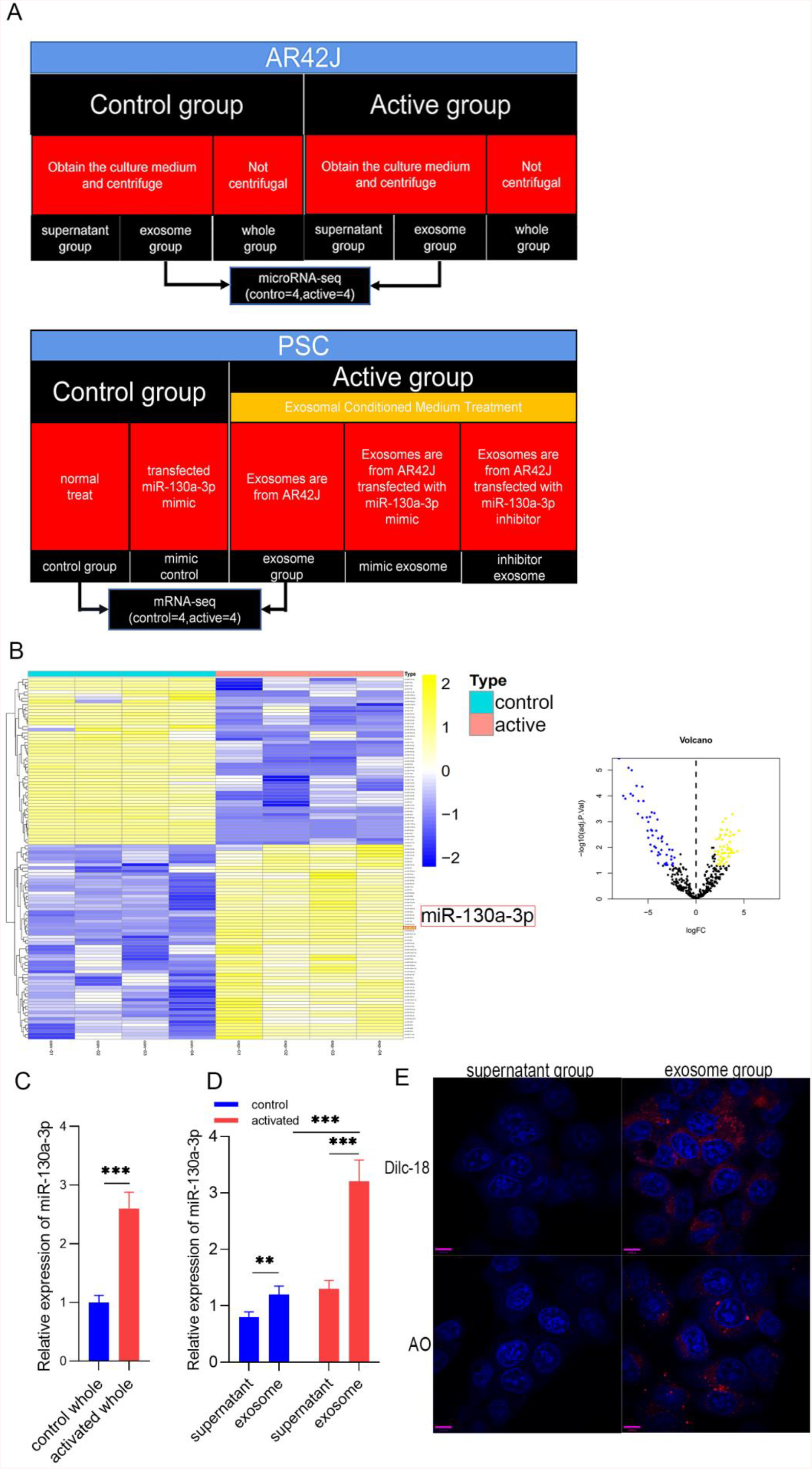
The expression, distribution and trajectory of exosomal miR-130a-3p. **A.** Cell experiment grouping situation. **B.** The expression of exosomal miRNA in the control group and in the activation group, in which miR-130a-3p expression is significantly high. **C** and **D**: Distribution of exosomal miR-130a-3p. **E** Dynamic tracing of exosomes and exosomal miRNAs. DilC-18 was used to label exosome phospholipid bilayers, and AO was used to label exosomal miRNAs. Red fluorescence represents exosomes and exosomal miRNAs, and blue fluorescence represents the PSC nucleus.

### PSC uptake of exosomes secreted by acinar cells and exosomal miRNA

To investigate the distribution of miR-130a-3p, we used RT-PCR to detect miR-130a-3p in the complete culture medium of control group and activated group cells. As expected, acinar cells released more miR-130a-3p after activation (Figure 1C). To further understand the distribution of miR-130a-3p, we centrifuged complete culture medium from the two groups to obtain exosomes and supernatant and then assessed the miR-130a-3p level in both. Surprisingly, we found that miR-130a-3p was mainly distributed in exosomes from both the control and activation groups, but the level of miR-130a-3p in the activated group was much higher than that in the control group. This result was consistent with our microarray analysis results (Figure 1D). To observe the trajectory of miRNAs in exosomes and exosomal miRNA, we labeled the miRNAs in exosomes and exosomes with fluorescent markers and observed them with a confocal fluorescence microscope. We observed no fluorescence in PSCs after addition of Dil-C18 labeled supernatant to PSC medium. In contrast, red fluorescence appeared in the PSCs (Figure 1E) after incubation with Dil-C18-labeled exosomes. This result confirmed that the exosomes were successfully extracted and were ingested by PSCs after incubation. AO was used to label miRNA in the supernatant group and the exosome group, which were then incubated with PSCs. In the AO-labeled supernatant group, no red fluorescence appeared in the PSCs, while in the AO-labeled exosomes group, red fluorescence appeared in the PSCs (Figure 1E). This result confirmed that miRNAs were abundantly enriched in the exosomes group and entered the PSCs through exosomes. In summary, we confirmed that miR-130a-3p was highly expressed in pancreatic acinar cells after injury and is mainly distributed in exosomes that transport miRNAs into PSCs.

### Exosomal miR-130a-3p derived from pancreatic acinar cells can activate PSCs and promote pancreatic fibrosis

To study the biological functions of acinar cell-derived exosomal miR-130a-3p after it enters PSCs, we extracted exosomes from acinar cells with overexpression or knockdown of miR-130a-3p, and incubated PSCs with the exosomes. RT-PCR was used to test the efficiency of cell transfection with miR-130a-3p. The results showed that the exosome mimic group had higher miR-130a-3p expression while the exosome inhibitor group had lower miR-130a-3p expression than the exosome group, indicating realization of in vitro control of miR-130a-3p expression (Figure 2A). We examined the expression of the PSC activation marker α-SMA protein and the fibrosis markers collagen I and collagen III at the RNA and protein level and found that the α-SMA, collagen I, and collagen III mRNA (Figure 2B-D) and protein (Figure 2E) levels in PSCs were significantly increased in the exosome group and mimic exosome group compared with the control group. Interestingly, the expression levels of α-SMA, collagen I, and collagen III in PSCs showed no differences at the mRNA level or protein level between the exosome group and the exosome mimic group. However, with knockdown of miR-130a-3p in the exosome inhibitor group, the α-SMA, collagen I, and collagen III mRNA (Figure 2B-D) and protein (Figure 2E) expression levels were significantly reduced and were close to normal levels in the PSCs. This result indicates that acinar cell-derived exosomal miR-130a-3p can activate PSCs and promote collagen overexpression by cells, leading to CP occurrence. However, this activation was only initiated with a certain amount of miR-130a-3p and did not rely on the level of expression. Furthermore, knockdown of exosomal miR-130a-3p significantly inhibited PSC activation and fibrosis (Figure 2B-D). More surprisingly, in the mimic control group, we directly overexpressed miR-130a-3p in PSCs without exosome involvement. Although miR-130a-3p expression was highest in this group (Figure 2A), the PSCs were not activated (Figure 2B) and did not show greater expression of fibrosis markers (Figure 2C-D). This result indicates that the activation of PSCs depends on exosomal miR-130a-3p derived from acinar cells.

**Figure 2.**
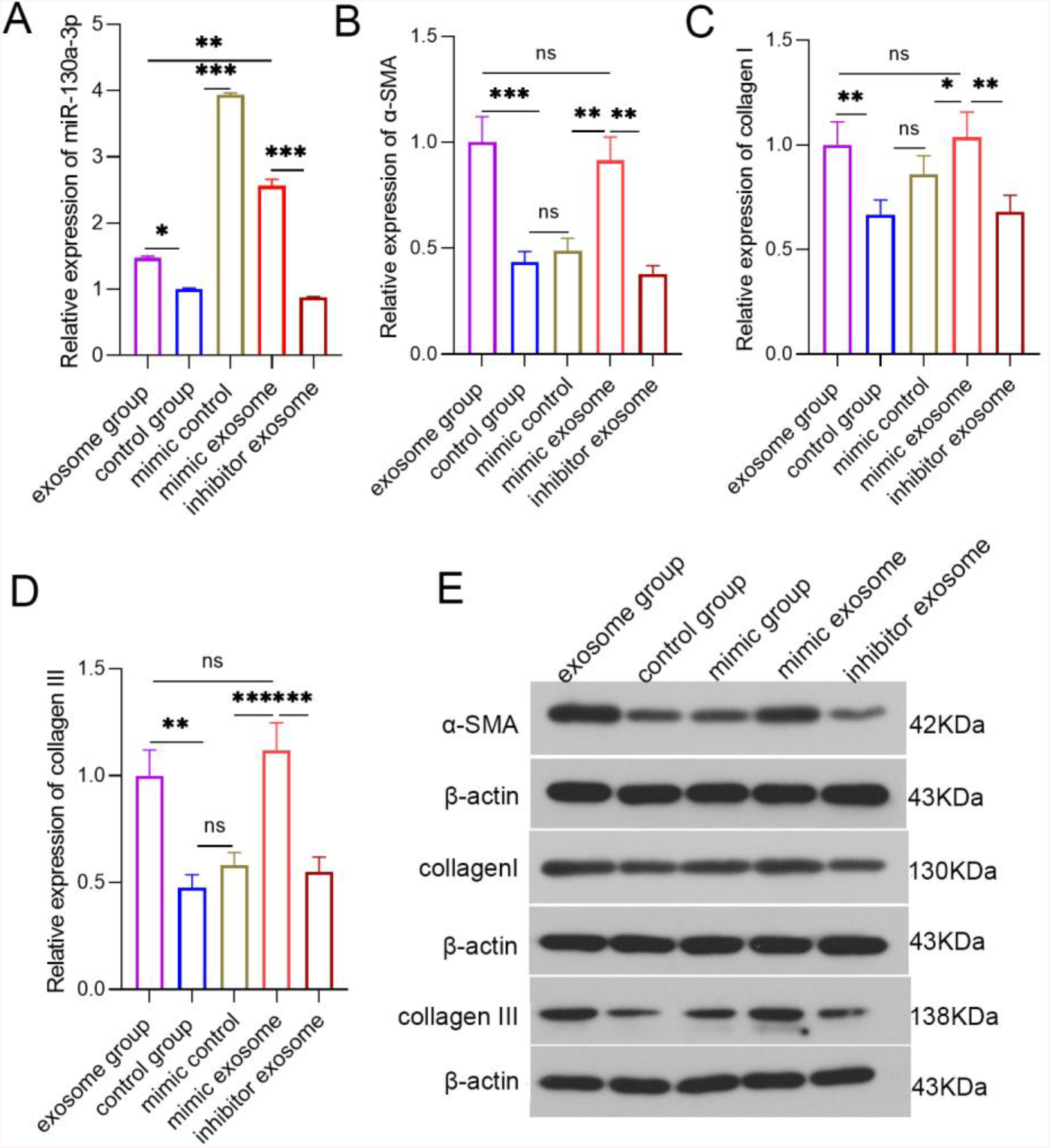
Detection of the various indicators in each group of PSCs at the cellular level via PCR or WB. **A** PCR detection of the relative expression level of miR-130a-3p in each group of PSCs. **B** PCR detection of α-SMA expression levels in each group of PSCs. **C** PCR detection of the collagen I expression level in each group of PSCs. **D** PCR detection of collagen III expression level in each group of PSCs. E WB detection of the α-SMA, collagen I and collagen III protein levels in PSCs.

To further explore the biological effects of miR-130a-3p in vivo, we divided rats into four groups: a sham operation group, CP group, inhibitor control group and inhibitor group. Among them, CP mice in the inhibitor group were transfected with miR-130a-3p inhibitor lentivirus in the pancreas. To test whether the model was successful, we used RT-PCR to detect miR-130a-3p expression in pancreatic tissues in each group. The results showed that miR-130a-3p expression in the CP group and the inhibitor control group was significantly higher than that in the inhibitor group or the sham operation group. The inhibitor group had the lowest miR-130a-3p expression. This result indicates that we achieved in vivo control of miR-130a-3p expression (Figure 3A). We examined the expression of the PSC activation marker α-SMA and the fibrosis markers collagen I and collagen III in pancreatic tissue and found that the expression of α-SMA, collagen I, collagen III mRNA (Figure 3B-D) and protein (Figure 3E) was significantly increased in the CP group versus the sham operation group. However, α-SMA, collagen I, and collagen III expression was significantly reduced in the inhibitor group versus the CP group (Figure 3B-E). H&E(Figure 4A)and Masson(Figure 4B)staining confirmed that the degree and area of fibrosis in the inhibitor group were significantly less than those in the CP group or the inhibitor control group (Figure 4D),indicating that pancreatic tissue fibrosis is associated with CP. Knockdown of miR-130a-3p in vivo significantly inhibited pancreatic tissue fibrosis. Immunohistochemistry also confirmed that knockdown of miR-130a-3p in vivo can inhibit the activation of PSC cells and reduce the expression of the specific protein α-SMA (Figure4C and E). In summary, PSCs are significantly activated at the occurrence of CP, and a significant increase in miR-130a-3p is associated with obvious pancreatic fibrosis. Moreover, knockdown of miR-130a-3p can inhibit PSC activation and prevent pancreatic fibrosis.

**Figure 3.**
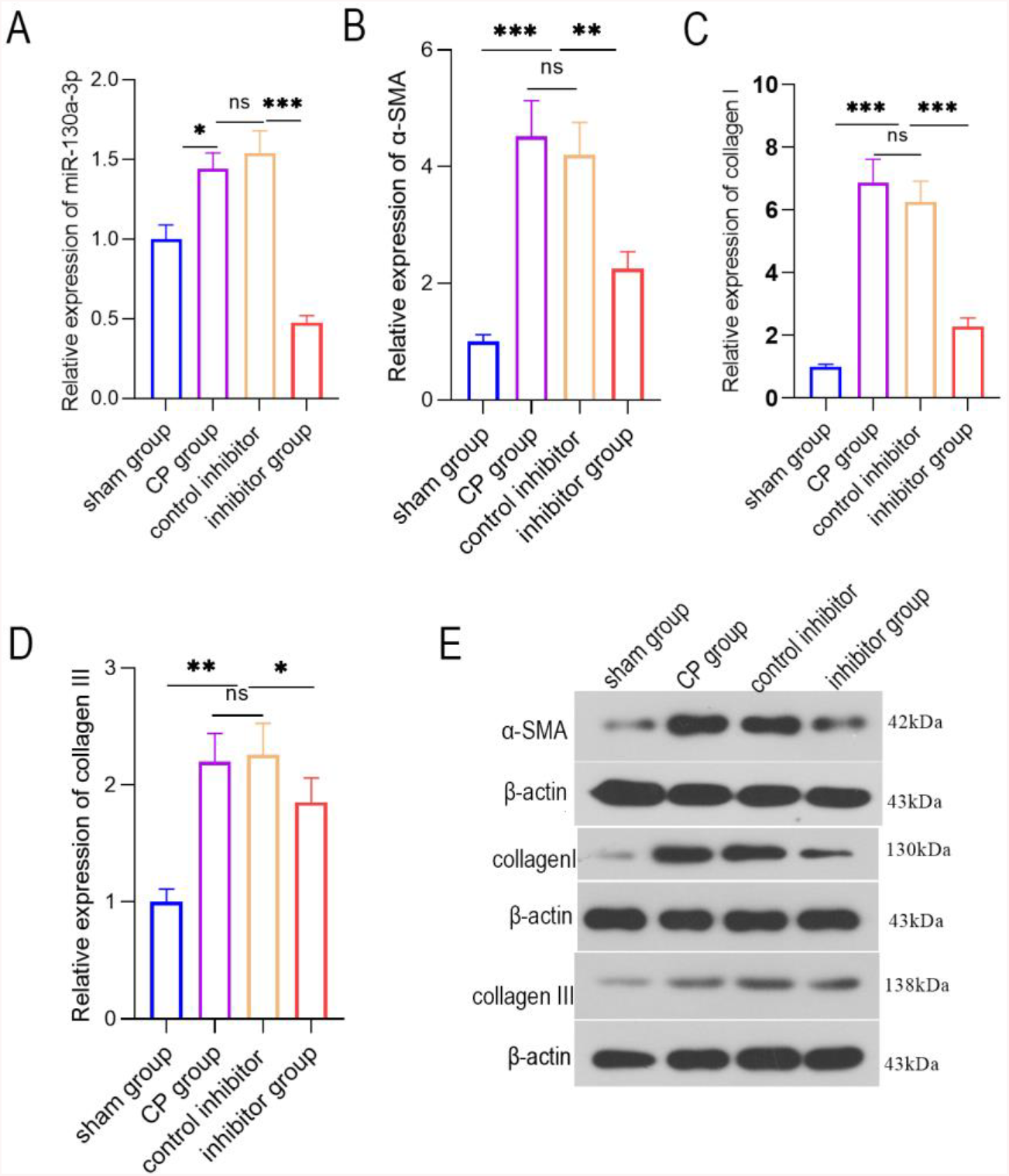
PCR or WB detection of the various indicators in pancreatic tissue from each group at the animal level. **A** PCR detection of the relative miR-130a-3p expression level in pancreatic tissue from each group. **B**PCR detection of the α-SMA expression level in pancreatic tissue from each group. **C** PCR detection of the collagen I expression level in pancreatic tissues from each group. **D** PCR detection of the collagen III expression level in pancreatic tissues from each group. **E** WB detection of the α-SMA, collagen I, and collagen III protein levels in pancreatic tissue.

**Figure4.**
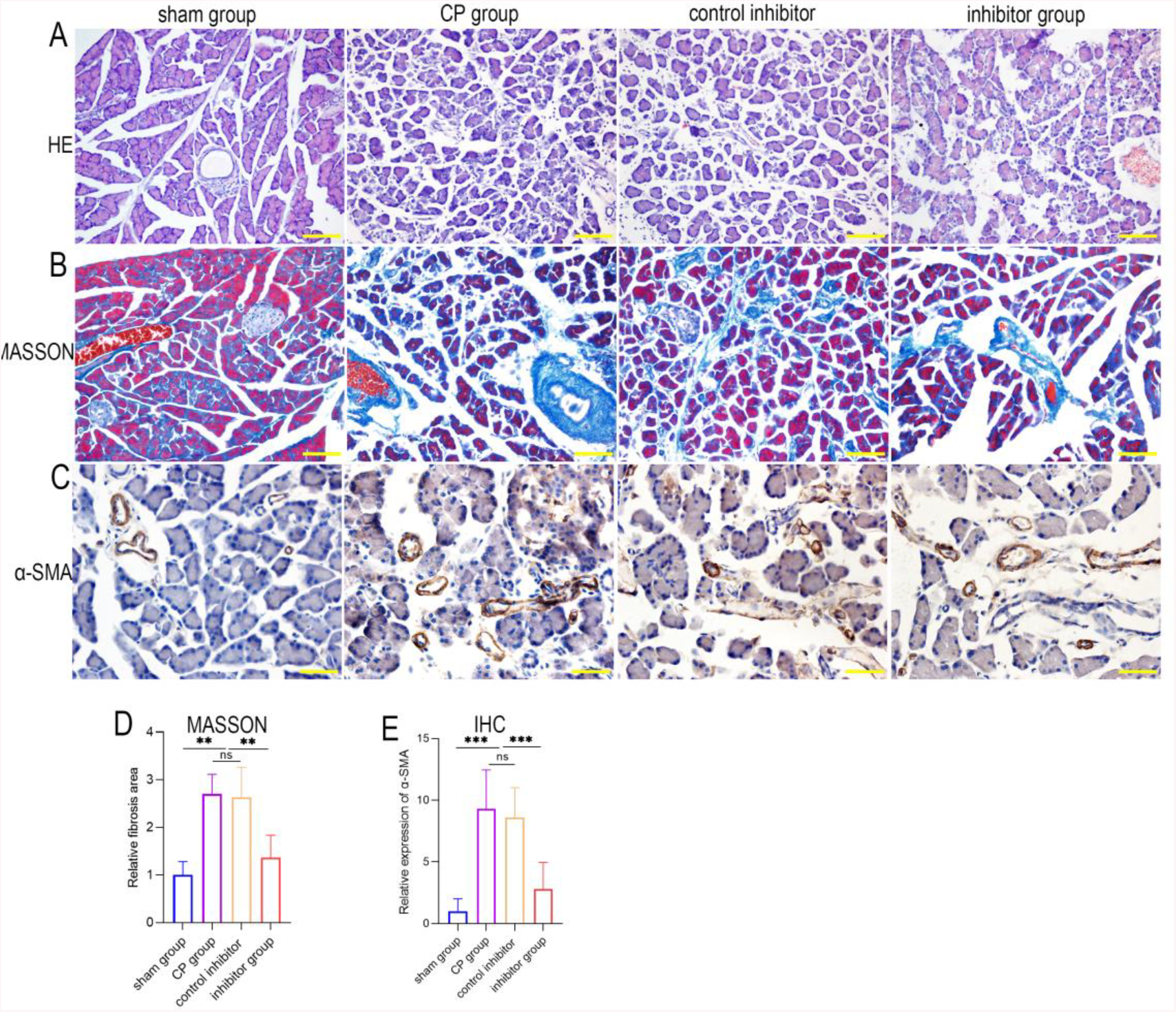
**A-B** H&E and Masson staining of pancreatic tissues from each group; **C** immunohistochemistry of α-SMA in pancreatic tissues from each group. **D** Masson of Fibrosis area in pancreatic tissues from each group. **E** Relative expression of α-SMA in pancreatic tissues from each group.

### Knockdown of miR-130a-3p not only improves pancreatic endocrine and exocrine functions but also protects endothelial cells

To investigate the role of miR-130a-3p in the recovery of the pancreas after pancreatic injury, we tested multiple indicators in the peripheral blood of four groups of animals. Of these indicators, C-peptide level reflects the endocrine function of the pancreas, with a lower level indicating worse function. The β-amylase level reflects the exocrine function of the pancreas, with a higher level indicating worse function. As expected, compared with the sham operation group, the CP group had a significantly lower C-peptide level and a significantly higher β-amylase level. This result indicates that the endocrine and exocrine functions of the CP group were severely damaged. However, surprisingly, knockdown of miR-130a-3p led to a significantly higher C-peptide level and a significantly lower β-amylase level in the inhibitor group than in the CP group (Figure 5A-B). Therefore, knockdown of miR-130a-3p significantly improved pancreatic endocrine and exocrine functions.

**Figure 5.**
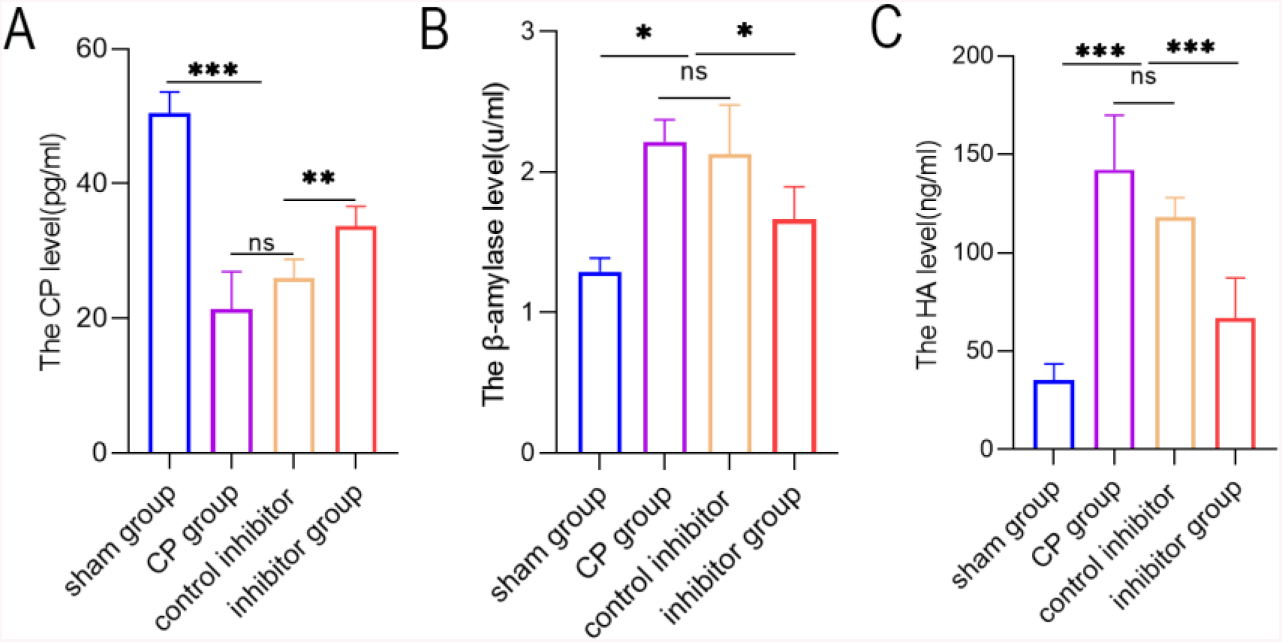
Knockdown of miR-130a-3p not only improves pancreatic endocrine and exocrine functions but also protects endothelial cells. **A** ELISA detection of C-peptide in serum from mice in each group. **B** ELISA detection of hyaluronic acid (HA) in serum from mice in each group. **C** DNS method for calculation of the β-amylase activity in serum from mice in each group.

Similarly, we tested the serum HA content in the four groups. The HA content reflects the function of endothelial cells, with a higher content indicating better function. We found that the HA content in the inhibitor group was significantly higher than in the CP group (Figure 5C). This result confirmed that knockdown of miR-130a-3p can effectively protect endothelial cell function. In summary, knockdown of miR-130a-3p can not only effectively improve pancreatic endocrine and exocrine functions but also protect endothelial cells and promote the recovery of pancreatic tissue.

### miR-130a-3p promotes pancreatic fibrosis by inhibiting PPAR-γ expression

In the early stage, we confirmed that miR-130a-3p is involved in the activation of PSCs and promotes pancreatic fibrosis. To investigate the underlying mechanisms and identify the downstream target genes of miR-130a-3p, we analyzed the differentially expressed mRNAs before and after PSC activation (Figure 6A) and constructed a PPI network (Figure 6B). We analyzed the centrality of the PPI network through cytoNCA and identified 10 essential proteins in the biological network (Figure 6B). Then, the TargetScan and miRanda databases were used to predict the target genes of miR-130a-3p and to find an intersection set of the target genes and the mRNA encoding the 10 essential proteins. Finally, we identified that PPAR-γ(PPARG) was significantly underexpressed. This low expression is consistent with the predicted trend of miRNA inhibition (Figure 6C). A literature review revealed that the PPAR-γ pathway has been confirmed to be involved in PSC activation and that PPAR-γ is a “star molecule”. Therefore, we performed an in-depth study of the miR-130a-3p-PPAR-γ pair. To verify that miR-130a-3p can bind to the target gene PPAR-γ and the regulatory relationship between them, we conducted a dual-luciferase reporter gene experiment. The results showed that the fluorescence activity difference between the WT-NC and MUT-NC group was not significant, but with overexpression of miR-130a-3p, the fluorescence activity in the WT+miR-130a-3p mimic group was significantly lower than that in the WT-NC group. This finding indicates that the expression of PPAR-γ was significantly suppressed. When the 3'UTR sequence of PPAR-γ was mutated, the fluorescence activity in the WUT+miR-130a-3p mimic group was restored to a higher level than that in the WT+miR-130a-3p mimic group, indicating that the expression of PPAR-γ was restored. This result confirms that the miR-130a-3p entering the PSCs can significantly inhibit PPAR-γ gene expression by directly binding to the PPAR-γ 3’UTR (Figure 6D).

**Figure 6.**
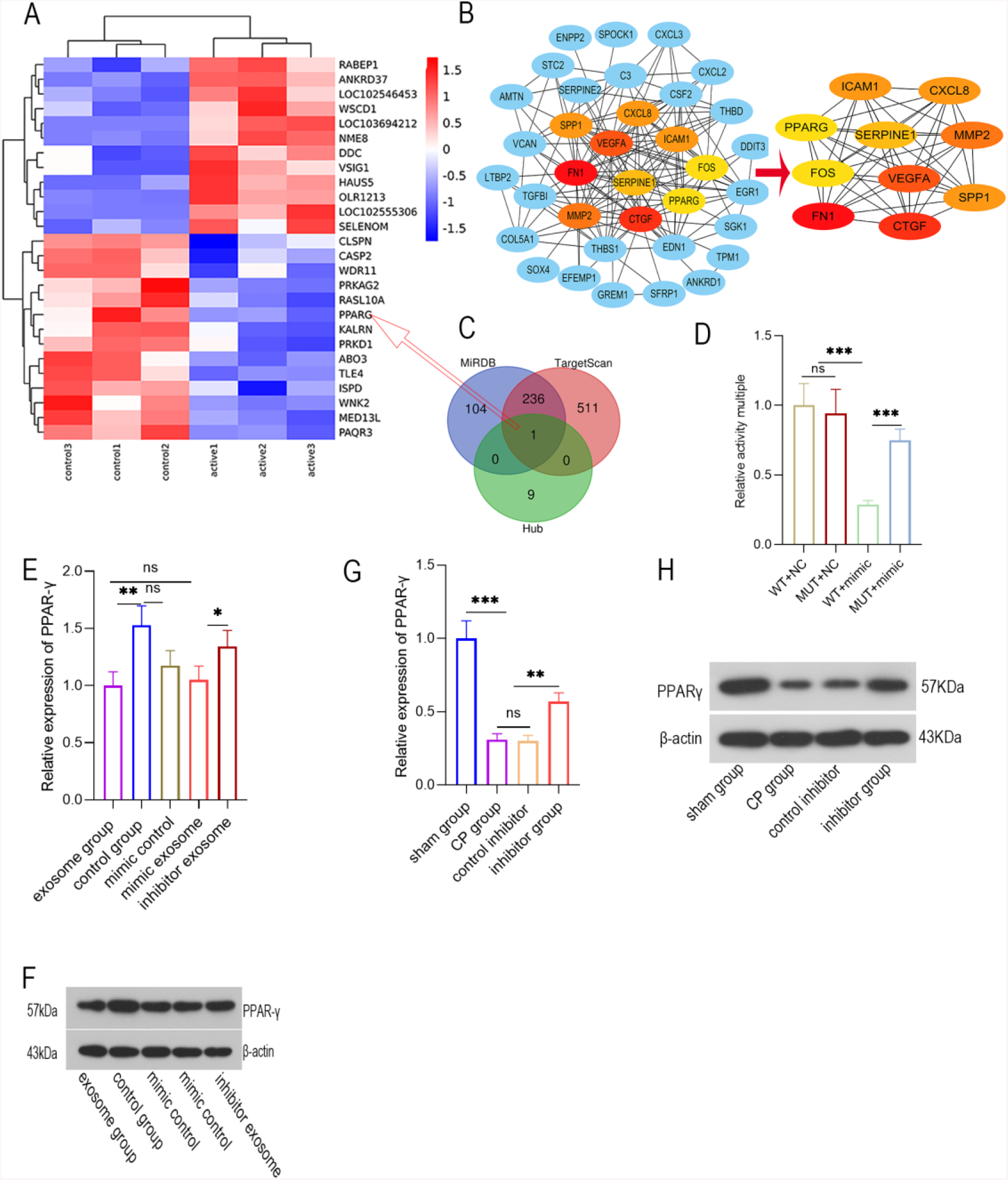
miR-130a-3p promotes pancreatic fibrosis by inhibiting PPAR-γ expression. **A** mRNA expression before and after PSC activation. **B** Based on the String database, a protein-protein interaction (PPI) network map was plotted for the significantly differentially expressed genes, and a centrality analysis of the PPI network through cytoNCA was conducted to obtain 10 essential proteins. **C** Venn diagram showing miR-130a-3p target genes and 10 essential protein mRNAs. **D** Dual-luciferase reporter gene assay for determination of relative PPAR-γ activity. **E and F** Detect the relative expression of PPAR-γmRNA and protein level in cells. **G** and **H** Detect the relative expression of PPAR-γ mRNA and protein level in the tissue.

To further verify the regulation of PPAR-γ by miR-130a-3p, exosomes were extracted from acinar cells with overexpression or knockdown of miR-130a-3p and then incubated with PSCs. Then, we examined the expression level of PPAR-γ. RT-PCR and western blotting showed that the expression of PPAR-γ was significantly lower in the exosomes group than in the control group, while PPAR-γ expression was increased in the exosomes group with knockdown of miR-130a-3p (Figure 6E-F). Once again, our results confirmed an inhibitory effect of exosomal miR-130a-3p on PPAR-γ expression. Subsequently, we further verified the regulation of PPAR-γ by miR-130a-3p at the animal level. The results showed that PPAR-γ expression in the CP group was significantly lower than in the sham operation group. However, when miR-130a-3p was knocked down, PPAR-γ expression was significantly increased. PPAR-γ is known to maintain the lipogenic properties of PSCs and maintain them in a quiescent state. Decreased expression of PPAR-γ can cause PSC activation, leading to pancreatic fibrosis; however, knockdown of miR-130a-3p can significantly upregulate PPAR-γ expression (Figure 6G-H).

## DISCUSSION

Pancreatic fibrosis is a chronic pathological process involving excessive accumulation of ECM in pancreatic tissue, and pancreatic mesenchymal PSCs play a key role in this process. PSCs are activated after being stimulated by a variety of factors, and the activated PSCs are transformed into myofibroblasts and migrate to the injured site to secrete excessive ECM, which eventually leads to pancreatic fibrosis. In this pathological process, exosomes and the miRNAs they carry play an important role. In the process of liver fibrosis, miR-181-5p carried by exosomes can regulate the STAT3/Bcl-2/Beclin 1 signaling pathway and reduce the expression of collagen fibers, thereby alleviating liver fibrosis^[14]^. In pancreatic cancer, cancer cells secrete a large number of exosomes, and the miRNA carried by them produces a fibrotic microenvironment that is conducive to cancer cell metastasis, thereby promoting liver metastasis of pancreatic cancer^[15,16]^. Masamune A et al. found that the exosomes secreted by pancreatic cancer cell lines (SUIT-2, Panc-1) contain a large amount of miR-1246 and miR-1290, which can stimulate PSC cell migration and activation and promote expression of α-SMA and fibrosis-related genes^[17]^. These studies have verified the important role of exosomes and the miRNAs they carry in the occurrence and development of diseases. In vitro fluorescence labeling and dynamic tracing showed that exosomes carry miRNAs into PSCs. Further studies confirmed that exosomal miR-130a-3p mediates information transmission between acinar cells and PSCs and activates PSCs, promoting pancreatic fibrosis. Previous studies have demonstrated that the miRNA-130 family is a key regulator of fibrosis pathways in 137 diseases and that inhibiting its expression can prevent ECM accumulation^[18]^. Gooch et al.^[19]^confirmed that miR-130 is involved in Cyclosporine A-mediated renal fibrosis. Chu et al. confirmed that miR-130 can promote NF kappa B-mediated inflammation and aggravate TGF-β1-mediated fibrosis by inhibiting the protective effect of the target gene PPAR-γ in a rat acute myocardial infarction model^[20]^. These findings suggest that miR-130 is involved in the pathological process of kidney^[21]^ and heart tissue fibrosis. How does it affect the process of fibrosis in pancreatic tissue? Our study shows that the occurrence of CP is associated with higher miR-130a-3p expression in acinar cells, which is mainly distributed in exosomes (Figure 1D). After incubation with PSCs, exosomes can carry miR-130a-3p into PSCs (Figure 1E) and stimulate PSC activation, leading to secretion of more collagen fibers. However, initiation of this activation only requires a certain amount of miR-130a-3p but does not rely on the level of miR-130a-3p expression. However, knockdown of exosomal miR-130a-3p can significantly inhibit PSC activation and inhibit fibrosis (Figure 2B-E, Figure 3 and 4). Notably, in the mimic control group, direct overexpression of miR-130a-3p in PSCs without exosome involvement did not activate PSCs or significantly increase ECM production. This finding indicates that PSC activation depends on acinar cell-derived exosomal miR-130a-3p and does not rely on exogenous miRNA (Figure 2B-D). In addition, knockdown of miRNA-130a-3p in animal models reduced the levels of serum HA and β-amylase but increased the C-peptide content (Figure 5A-C). This result indicates that knockdown of miR-130a-3p can not only inhibit pancreatic fibrosis but also protect pancreatic endocrine, exocrine and endothelial cell functions.

In this regard, which molecules act as a target of miR-130a-3p? To identify this pathway, we conducted a series of bioinformatics analyses and determined from the entire network that PPAR-γ is a target of miR-130a-3p (Figure 6B-C). PPAR-γ plays an important role in maintaining PSC quiescence. PPAR-γ can maintain the adipogenic properties of PSCs and inhibit PSC proliferation, collagen synthesis and α-SMA expression^[22]^. High PPAR-γ expression can inhibit PSC activation and the promotion of fibrosis induced by PDGF and thus has potential application value in the treatment of CP^[23,24]^. The PPAR-γ receptor agonist troglitazone inhibits PSC proliferation, collagen synthesis and α-SMA expression by blocking G1 phase of the PSC cell cycle and reduces CP fibrosis^[25,26]^. These previous studies have shown that PPAR-γ is a key molecule in pancreatic fibrosis^[27,28]^. However, PPAR-γ is significantly underexpressed in CP (Figure 6A). How is it regulated? Through a dual-luciferase gene report experiment, we confirmed that exosomal miRNA-130a-3p can directly bind to PPAR-γ and inhibit its expression (Figure 6D). To verify this conclusion, we performed experiments to overexpress miRNA-130a-3p in vitro and in vivo, and the results showed that PPAR-γ expression decreased at both the transcriptional and protein levels and that pancreatic fibrosis was aggravated compared with the control group. In the case of miRNA-130a-3p knockdown, PPAR-γ expression significantly increased at both the transcriptional and protein level, and the degree of fibrosis significantly improved (Figure 6E-H). Therefore, the miRNA-130a/PPAR-γ regulatory axis may become a new target to regulate PSCs against pancreatic fibrosis.

## CONCLUSIONS

In summary, this study demonstrated that damaged pancreatic acinar cells release exosomes that carry miR-130a-3p and transport miR-130a-3p to PSCs. By inhibiting PPAR-γ expression, miR-130a-p promotes PSC activation and collagen formation, leading to fibrosis of pancreatic tissue, whereas knockdown of miR-130a-3p can significantly inhibit pancreatic fibrosis. CP fibrosis is an important pathological feature of pancreatic cancer^[29;30]^. Investigating the mechanism of pancreatic fibrosis has far-reaching clinical significance. This study provides new ideas for the prevention and treatment of CP fibrosis from the perspective of exosome-mediated intercellular communication.

## METHODS

### Ethics statement

The animals were managed in accordance with a protocol approved by the local animal use and care committee and were euthanized in accordance with the National Animal Welfare Law

### Cell culture

Rat pancreatic acinar cells (AR42J, The Cell Bank of Type Culture Collection of Chinese Academy of Sciences) were cultured in Ham’s F12K medium containing 20% fetal bovine serum, 100 kU/L penicillin and 100 mg/L streptomycin in a 37 °C and 5% CO_2_ incubator. The PSCs were obtained from clean-grade healthy male SD rats weighing 200-300 g. The cells were inoculated in culture flasks for further culture. PSC identification was performed every day.

Treatment of rat pancreatic acinar cells (Figure 1A):The cells in the control group were routinely cultured in Dulbecco’s modified Eagle’s medium (DMEM), while the cells in the activation group were cultured in DMEM containing 200 μg taurolithocholate TLC-S for 48 h,then cells were separated and washed with PBS. Then, the culture medium was replaced with fresh medium, and the cell culture continued for another 48 h. The culture medium was collected and centrifuged. The supernatant was obtained and retained as the supernatant group. The white precipitate in the bottom of the vial was resuspended in DMEM and saved as the exosome group (a part of this suspension was saved as conditioned medium for future use and the other part was used for extraction of exosomes and their miRNAs). Cell culture medium that was not centrifuged was used as the complete culture medium group. In total, six groups were examined in this experiment: control exosome group, control supernatant group, control whole culture medium group, activated exosome group, activated supernatant group, and activated whole culture medium group.

Treatment of rat pancreatic stellate cells (Figure 1A): The exosomes were incubated with rat PSC for 14 days, and then the PSC was subjected to mRNA-seq.The PSCs were divided into five groups. In the control group, PSCs were treated with normal culture medium for 14 days. In the exosome group, rat PSCs were treated with conditioned medium for 14 days. In the mimic control group, PSCs in the control group were transfected with miR-130a-3p mimic on the 11th day. In the mimic exosomes group, PSCs were incubated for 14 days with exosomes extracted from AR42J cells transfected with miR-130a mimic. In the inhibitor exosomes group, PSCs were incubated for 14 days with exosomes extracted from AR42J cells transfected with miR-130a inhibitor. Transfection was performed using LipofectamineTM2000 (Invitrogen, Carlsbad, CA) according to the manufacturer’s instructions. The miR-130a-3p mimic sequence was CAGUGCAAUGUUAAAAGGGCAU, and the rno-miR-130a inhibitor sequence was AUGCCCUUUUAACAUUGCACUG.

### Animals

The Animal Research Center of the First Clinical College of Harbin Medical University provided 20 healthy male SD rats (250±20 g). The rats were randomly divided into four groups and allowed to acclimate for one week. The rats were fasted with free access to water for 12 h before surgery. A midline incision was made along the linea alba, and the pancreatic duct was exposed at the junction between the stomach and the duodenum. In the sham operation group, ligation of the pancreatic duct was not performed, and an equal volume of sterile saline was injected intraperitoneally. In the CP group, ligation of the pancreatic duct was performed with intraperitoneal injection of 50 μg/kg cerulenin two days later^[8]^. In the negative interference lentivirus group, 100 μl of lentivirus with random interference sequences was injected into the biliary and pancreatic duct in animals from the model group. In the inhibitor group, 100 μl of miR-130a inhibitor lentivirus (1×10^8^ pfu/ml) was injected into the biliary and pancreatic duct in animals from the model group^[9]^. The skin incision was closed in layers. One month later, the animals were euthanized after blood was sampled from the inferior vena cava. Parts of the pancreatic tissues were collected and fixed in 10% neutral formaldehyde solution and 2.5% glutaraldehyde.

### Extraction of exosomes and exosome miRNAs from rat pancreatic acinar cell culture medium

We used ExoQuickTM exosome precipitation solution (SBI, USA) for exosome extraction. TRIzol reagent was used to directly extract total RNA, including miRNA, from exosomes.

### Microarray-based gene expression profiling

MiRNAs:After qualified quality control of total RNA was confirmed, a microarray analysis of miRNAs was performed, including labeling, hybridization, signal amplification and image acquisition. Then, data were extracted from the obtained images and analyzed. mRNA: The Agilent SurePrint G3 Rat Gene Expression v2 8×60K Microarray(DesignID:074036) was used in this experiment and data analysis of the 6 samples wereconducted by OE Biotechnology Co., Ltd., (Shanghai, China). Total RNA was quantified by the NanoDrop ND-2000 (Thermo Scientific)and the RNA integritywas assessed using Agilent Bioanalyzer 2100 (Agilent Technologies).The sample labeling,microarray hybridization and washing were performed based on the manufacturer’s standardprotocols. Briefly, total RNA were transcribed to double strand cDNA, then synthesizedinto cRNA and labeled with Cyanine-3-CTP. The labeled cRNAs were hybridized onto the microarray.After washing, the arrays were scanned by the Agilent Scanner G2505C (Agilent Technologies).

### RT-PCR quantitation

According to the instructions of an RNA extraction kit (BioTeke, RP1201, China), total RNA was extracted, cDNA was synthesized by reverse transcription, and real-time fluorescence quantitative analysis was performed on an ExicyclerTM 96 fluorescence detector. All data were analyzed via relative quantitative analysis using the 2^-ΔΔCt^ method to detect the levels of PPAR-γ, α-SMA, Collagen I and Collagen III mRNAs. The experiment was repeated three times.

### Dynamic tracing of exosomes and exosomal RNAs

AR42J cells (7×10^6^) were plated. pPACKH1 (45 μL) and CD9 Cyto-TraceTM plasmid (4.5 μg) were added into fresh culture medium with PureFectionTM (5.5 μL). The medium was mixed for 10 sec, kept at room temperature for 15 min and then added into the culture medium of AR42J cell. The cells were cultured in a 37 °C and 5% CO_2_ incubator for 24 h. The AR42J cells in the culture medium were treated with Dil-C18 fluorescent dye for 1 h and then washed with PBS three times to label the phospholipids on the exosome membrane, and the exosomes were identified with a laser confocal microscope.

The principle of Exo-RedTM staining is based on the properties of Acridine Orange (AO). AO can pass through the cell membrane and fluorescently label single-stranded RNAs inside the exosomes. The detailed protocol was as follows: the obtained exosomes were resuspended in 500 μL 1×PBS in a 1.5 mL EP tube; 50 μL of 10×Exo-RedTM was added to the resuspended exosome solution; the mixture was incubated at 37 °C for 10 min; 100 μL of ExoQuick - TCTM reagent was added to stop the labeling reaction; the mixture was placed on ice for 30 min and then centrifuged at 14,000×g for 3 min; the supernatant was removed; the labeled exosomes were resuspended in 500 μL of 1×PBS; 100 μL of labeled exosomes were added to each well of a six-well plate (approximately 1×10^5^ PSCs per well); the cells were observed using a confocal laser microscope after 24 h of culture.

### Western blot analysis

A kit (Wanleibio, WLA019) was used to detect the PPAR-γ, α-SMA, Collagen I and Collagen III levels, with β-actin antibody as an internal control.

### ELISA and DNS

Serum was obtained by centrifugation of the inferior vena cava blood of rats in the four groups. β-Amylase activity was calculated using a β-amylase detection kit (Nanjing Jianshe, C016-2) and the DNS method. A hyaluronic acid (HA) detection kit (Ulsan, CEA182Ge) and C-peptide detection kit (Ulsan, CEA447Ra) were used to calculate the concentration of HA and CP based on the ELISA method according to the manufacturer’s instructions.

### H & E staining, Masson staining and immunohistochemistry

H & E staining and Masson staining were performed on the pancreatic tissues of rats in the four groups. After staining, the stained sections were photographed under a microscope at 200× magnification. Immunohistochemistry for α-SMA was performed, and the sections were photographed under a microscope at 400× magnification. HistoQuest software was used for quantitative analysis of Masson and immunohistochemical staining intensity.

### Construction of miRNA-mRNA pairs

Data processing of the Exosomal miRNA expression profile chip and analysis of differentially expressed genes(control=4,active=4): the limma package in Bioconductor was used to analyze differentially expressed genes. The settings false discovery rate (FDR) <0.05 and |log2FC|≥0.58 (that is, the expression difference between groups was not less than 1.5 fold) were chosen. Data processing of the PSC mRNA expression profile chip and analysis of differentially expressed genes(control=3,active=3): the limma package in Bioconductor was used to analyze differentially expressed genes. The settings false discovery rate (FDR) <0.05 and |log2FC|≥0.58 (that is, the expression difference between groups was not less than 1.5 fold) were chosen. Based on the String database, a protein-protein interaction (PPI) network was constructed for significantly differentially expressed mRNA (the threshold was a PPI confidence score=0.4). Then, we used cytoNCA^[10,11]^to analyze the centrality of the PPI network. We used eight centrality measures (Betweenness, Closeness Centrality, Degree Centrality, Eigenvector Centrality, Local Average Connectivity-based Centrality, Network Centrality, Subgraph Centrality, Information Centrality) to identify 10 essential proteins in the biological network. Then, the TargetScan and miRanda databases were used to predict the target genes of miR-130a-3p. The intersection of the target gene and the mRNA encoding 10 essential proteins was obtained to construct the miRNA-mRNA pairs.

### Determination of luciferase reporter gene

The miRDB^[12]^database and DIANA TOOLS^[13]^ database were used to predict the binding site of miR-130a-3p to the PPAR-γ gene. In the putative binding sites, the 3’UTR sequence of the PPAR-γ gene that contains 3 bp mutation, the 3’UTR sequence of PPAR-γ and its mutant were inserted separately into the luciferase reporter plasmid psiCHECK-2 to construct wild-type (PPAR-WT) and mutant (PPAR-MIT) recombinant plasmids. Next, PSCs were transfected with miR-130a-3p mimic or negative control (NC) and the above-depicted recombinant plasmids. Forty-eight hours after transfection, a dual-luciferase detection kit was used to detect luciferase activity. The PSCs were divided into four groups: WT+NC, MUT+NC, Wt+mimic, and MUT+mimic groups.

### Statistical analysis

SPSS 20.0 software was used for statistical analysis, and the experimental data are all expressed as the mean ± standard deviation (x ± s). A t test was used to compare two sample means, and one-way analysis of variance (one-way ANOVA) was used to compare the means between multiple groups.

## ABBREVIATION

TLC: taurolithocholate;
PSC: pancreatic stellate cell;
HA: hyaluronic acid;
CP: Chronic pancreatitis;
ECM: extracellular matrix;
PSCs: pancreatic stellate cells;
AP: acute pancreatitis;
miRNAs: microRNAs;
DMEM: Dulbecco’s modified Eagle’s medium;
AO: Acridine Orange
HA: hyaluronic acid;
FDR: false discovery rate;
PPI: protein-protein interaction;

## DECLARATION

## ACKNOWLEDGMENTS

The authors would like to acknowledge the helpful comments on this paper received from the reviewers

## AUTHOR CONTRIBUTIONS

Dongbo Xue,Weihui Zhang and Chenjun Hao conceived the study;Qiang W ang and Hao Wang designed and performed the experiments;Qingxu Jing,Yang Yang performed experiments;Qiang Wang,Hao Wang performed the in vivo experiments;Qingxu Jing,Yang Yang performed the in vitro experiments;Qiang Wang conducted a series of bioinformatics analysis; All authors revised the manuscript.

## ETHICS APPROVAL AND CONSENT TO PARTICIPATE

The experimental design was approved by the Ethics Committee and Experimental Animal Ethics Committee of The First Affiliated Hospital of Harbin Medical University (ChiCTR1900024765).All animal experiments were performed in strict agreement with the guidelines of animal care and use. Extensive efforts were made to ensure minimal suffering of the animals used in the study.

## CONSENT FOR PUBLICATION

Not applicable.

## AVAILABILITY OF DATA AND MATERIAL

The datasets generated/analyzed during the current study are available.

